# Interpretable Predictive Modeling for Medical Data Using Boolean Rule-aware Regression

**DOI:** 10.64898/2026.05.14.725084

**Authors:** Mohammad Eskandarian, Seyed Amir Malekpour

**Affiliations:** Faculty of Mathematics and Computer Science, TU Bergakademie Freiberg (TUBAF), 09599, Freiberg, Germany; School of Biological Sciences, Institute for Research in Fundamental Sciences (IPM), Shahid Lavasani, Tehran, 19395-5746, Tehran, Iran

**Keywords:** Explainable artificial intelligence, Rule based systems, Ridge regression, Bayesian Information Criterion

## Abstract

**Purpose:** In clinical practice, accurate prediction of disease risk must be accompanied by transparent, human-understandable explanations to support diagnostic confidence, guide therapeutic decisions, and meet ethical and regulatory standards. While deep neural networks achieve high predictive performance in tasks such as cancer detection and diabetes risk stratification, their black-box nature prevents clinicians from understanding the reasoning behind predictions, severely limiting trust and safe integration into patient care.

**Methods:** We present Regression-Based Boolean Rule (RBBR), a framework that automatically derives clinically interpretable Boolean rules directly from patient data. RBBR generates human-readable conjunctions (logical AND combinations) of up to three clinical features, transforms them into inputs for ridge regression to predict binary or multi-class disease outcomes, estimates rule importance via regularized coefficients, and selects the most parsimonious and predictive rule sets using the Bayesian Information Criterion.

**Results:** Applied to six real-world medical datasets (lung cancer screening and staging, Wisconsin and diagnostic breast cancer, heart failure, and early-stage diabetes risk), RBBR consistently produced concise, clinically meaningful rules – e.g., gender-specific symptom combinations in diabetes, distinct histopathological subpopulations in breast cancer, and symptom-risk factor interactions in lung cancer – with strong explanatory power (***R***^***2***^ up to 0.92) and competitive discrimination.

**Conclusion:** By delivering logical, transparent decision rules aligned with clinical reasoning (if symptom A **and** B, then high risk), RBBR bridges the gap between predictive accuracy and bedside usability, enabling clinicians to validate predictions, identify high-risk patients, stratify subpopulations, and enhance shared decision-making in routine care.

## Introduction

In medical data analysis, the tension between predictive accuracy and model interpretability is a persistent challenge. Predictive accuracy, is essential for reliable clinical prediction, including disease presence or risk estimation. However, interpretability, which ensures that a model provides transparent, human-understandable explanations, is equally critical in healthcare, where clinicians need insight into decision rationales to trust and act on predictions [1]. Interpretability reveals which features drive predictions and why, fostering trust and facilitating integration into clinical workflows.

Deep neural networks (DNNs) exemplify this trade-off, achieving high predictive accuracy in tasks such as breast cancer image classification or diabetes risk prediction [2]. However, their complexity, characterized by multilayer architectures and millions of parameters, makes them inherently opaque, often described as black boxes due to their lack of transparency [3]. This opacity, stemming from nested input, hidden, and output layers, obscures feature interactions, making it difficult to trace how inputs, such as patient symptoms, lead to outputs, such as disease risk, without post-hoc explanations. While DNNs offer benefits like scalability and accuracy, their lack of interpretability hinders clinical adoption, as clinicians cannot validate predictions without understanding the underlying logic [4]. Furthermore, integrating rule-based optimization into DNNs is computationally intractable, limiting their transparency even further [5, 6]. This lack of transparency is particularly problematic in high-stakes decision-making scenarios, such as medical diagnostics, where trust and patient safety are paramount. Clinicians need to understand why a model predicts heart failure based on exercise test results or flags diabetes risk from symptoms like polyuria [7]. Without this insight, black-box models risk misdiagnosis or missed interventions, as their predictions may reflect spurious correlations rather than underlying biological mechanisms. In such contexts, explainability is not merely desirable but necessary for ethical and regulatory compliance [8].

Building upon our recent work [9, 10], we propose a Regression-Based Boolean Rule (RBBR) framework for inferring Boolean rules from medical data. RBBR transforms input features into human-readable conjunctions (Boolean rules), such as “if *Sex*∧*ChestPainType*∧*Oldpeak*, then heart disease risk” where ∧ denotes the Boolean AND operator. We consider conjunctions or Boolean rules involving single, double, and triple-feature combinations [11, 12]. This choice is guided by previous research, which suggests that using up to three features in a rule provides an optimal balance between complexity and predictive accuracy [13, 14]. To assess the predictive power of these Boolean rules, we use them as inputs in a ridge regression model, where each rule is assigned a coefficient reflecting its relevance and efficacy in explaining the target class, such as disease status or diagnosis. We use ridge regression to effectively handle multicollinearity among Boolean rules [15] and ensure robust performance on sparse datasets [16]. By providing each ridge regression model with a set of Boolean rules as inputs for predicting the target class, we identify the optimal rules with the highest coefficients, highlighting their predictive power. For example, consider a Boolean rule set { (*Age* ∧ *Alcohol*), (*Age* ∧ ¬*Alcohol*),(¬*Age* ∧ Alcohol), (¬*Age* ∧ ¬*Alcohol*)} involving two features, *Age* and *Alcohol*, (¬ represents negation). When fitting a ridge regression model with these rules as inputs, only the rule (*Age* ∧ ¬*Alcohol*) may receive a high coefficient. This suggests that the presence of the feature Age, even in the absence of Alcohol, is strongly associated with the disease. To maximize interpretability without compromising accuracy, RBBR employs Bayesian model selection via the Bayesian Information Criterion (BIC) to rank ridge regression models fitted to different Boolean rule sets. The Boolean rules from models with optimal BIC can then be applied for disease status or diagnosis prediction in medical studies. Unlike DNNs, RBBR automatically selects the most predictive rule sets and their associated risk factors or features from both binary and continuous features. With ∨ as the Boolean OR operator, for example, in the *Wisconsin Breast Cancer* dataset, RBBR inferred the Boolean rule set (*Cl*.*thickness* ∧ ¬*Cell*.*size* ∧ *Bare*.*nuclei*)∨(*Cl*.*thickness* ∧ *Cell*.*size* ∧ ¬*Bare*.*nuclei*) with an *R*^2^ of 0.87 and a BIC of −2425, demonstrating its explanatory power and model fit. This rule set suggests two distinct tumor subpopulations, where malignancy is indicated by clump thickness combined with either a large cell size or bare nuclei – two opposing features. See the text for further analysis. In diabetes prediction, RBBR infers the Boolean rule set (*Gender* ∧ *Polyuria* ∧ *Polydipsia*)∨(¬*Gender* ∧ *Polyuria* ∧ *Polydipsia*)∨(¬*Gender*∧ ¬*Polyuria* ∧ *Polydipsia*)∨(¬*Gender* ∧ *Polyuria* ∧ ¬*Polydipsia*) which captures classic hyperglycemia symptoms while reflecting gender-specific physiological variations. In males, the rule (*Gender* ∧ *Polyuria* ∧ *Polydipsia*) suggests that both polyuria and polydipsia are necessary indicators of diabetes risk. In contrast, for females, the remaining rules indicate that the presence of either symptom alone may be sufficient to signal risk.

RBBR effectively captures complex feature interactions through Boolean rules, making it well-suited for medical applications where both accuracy and interpretability are essential. By deriving interpretable rules, RBBR enhances clinical trust and usability. These rules naturally align with medical decision-making, where diagnoses often depend on logical combinations of symptoms or test results, and can be relevant to distinct risk groups or subpopulations within the studied samples. Its application across datasets such as lung and breast cancer, heart disease, and diabetes demonstrates RBBR’s potential to support clinical decision-making and establish a foundation for eXplainable Artificial Intelligence (XAI) in healthcare.

## Materials and Methods

In what follows, we detail the RBBR framework: (i) generating Boolean rules (conjunctions). In high-dimensional datasets, to strike the right balance between accuracy and interpretability, we use Boolean rules involving up to three input features to explain the target class [13, 14], (ii) fitting a ridge regression model using a repertoire of Boolean rules as inputs and the target class as the response, (iii) deriving the optimal Boolean rules that best explain the target class (e.g., those with the highest coefficients in the fitted ridge regression model), and (iv) utilizing the Bayesian Information Criterion (BIC), we rank distinct ridge regression models fitted to Boolean rule sets, each involving different subsets of input features.

### Dataset Description

The RBBR framework was evaluated on six medical datasets, including Lung Cancer Prediction datasets with binary and three-level target class, Wisconsin Breast Cancer, Diagnostic Breast Cancer, Heart Failure Prediction, and Early Stage Diabetes Risk. These datasets vary in sample size, feature count, and target class distribution, reflecting diverse clinical contexts. The key characteristics of these datasets are summarized in Table 1, and they are available from Kaggle and UCI Machine Learning Repository[17].

**Table 1.** Characteristics of the Utilized Medical Datasets.

| Dataset (Source) | # Samples | # Features | # Classes | Majority Class (%) |
| --- | --- | --- | --- | --- |
| Lung Cancer Prediction (Kaggle) | 59 | 4 | 2 | 52.5 |
| Three-level Lung Cancer (Kaggle) | 1000 | 23 | 3 | 36.5 |
| Wisconsin Breast Cancer (UCI) | 699 | 9 | 2 | 65.5 |
| Diagnostic Breast Cancer (UCI) | 569 | 30 | 2 | 62.7 |
| Heart Failure Prediction (Kaggle) | 746 | 11 | 2 | 52.3 |
| Early Stage Diabetes Risk (UCI) | 520 | 16 | 2 | 61.5 |

### Boolean Rule Generation

RBBR transforms clinical features {*A, B*, …, *L*} – such as *Gender, Polyuria*, or *Polydipsia* – into human-readable Boolean rules or conjunctions. For example, from a two-feature subset {*A, B*}, it is possible to construct the following Boolean rule set consisting of four rules:

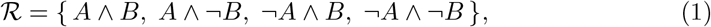

where ∧ denotes the Boolean AND operator and ¬ represents negation. For higherorder combinations, a three-feature subset {*A, B, C*} generates 2^3^ = 8 rules (e.g., *A*∧*B*∧*C, A*∧*B*∧¬*C*, …, ¬*A*∧¬*B*∧¬*C*). We construct Boolean rules using subsets of up to three clinical features [18, 13, 14]. This approach ensures that Boolean rules, such as (*Gender* ∧ *Polyuria* ∧ *Polydipsia*), remain interpretable and actionable for clinicians while capturing essential feature interactions. In practice, clinicians can use these rules to identify high-risk patients and make informed decisions, such as recommending lifestyle changes, further testing, or treatment. To compute the rule value for each sample, whether for training or prediction: (i) continuous or multi-level discrete features are first rescaled to the (0, 1) interval, and (ii) rule values are calculated consistently with prior studies on binary features by multiplying the corresponding feature values. For example, for the rule (*Gender* ∧ *Polyuria* ∧ *Polydipsia*), the output is computed as *Gender* × *Polyuria* × *Polydipsia*, while for (¬*Gender* ∧ ¬*Polyuria* ∧ *Polydipsia*), it is (1 − *Gender*) × (1 − *Polyuria*) × *Polydipsia*.

### Ridge Regression Formulation

The Boolean rules serve as input variables to a ridge regression model, predicting patient outcomes (e.g., disease risk). The model is formulated as:

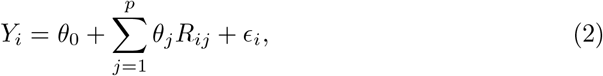

where *Y*_*i*_ denotes the outcome, and *R*_*ij*_ ∈ ℛ represents the *j*-th Boolean rule for the *i*-th patient. For instance, for the *i*-th patient, the values of the *p* = 4 Boolean rules involving input features *A* and *B* in (1) are given as follows:

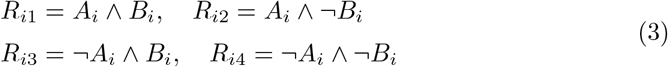

where *i* = 1, …, *n*, and *n* is the number of patients. In (2), *θ*_0_ is the intercept, *θ*_*j*_ is the regression coefficient for the *j*-th Boolean rule (*j* = 1, …, *p*), and *p* is the total number of Boolean rules. Additionally, _*i*_ represents the residual error, capturing noise or other small variations in patient outcomes that cannot be explained by the Boolean rules. In the presence of multicollinearity among predictors (Boolean rules) in (2), we use ridge regression to obtain stable and reliable parameter estimates. Ridge regression is a regularization technique that improves the stability and performance of linear regression models by adding a penalty term to the loss function – specifically, the squared magnitude of the coefficients scaled by a tuning parameter *λ*. This approach mitigates multicollinearity and prevents overfitting, particularly in noisy medical datasets. The objective function for parameter estimation in ridge regression is:

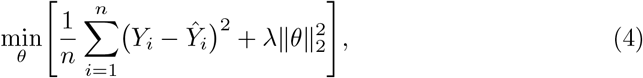

where 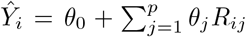, and 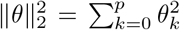 is the 𝓁_2_-norm of the coefficient vector. Additionally, *λ* is the regularization parameter tuned via 5-fold cross-validation to optimize the bias-variance trade-off [19].

### Bayesian Information Criterion (BIC)

RBBR explores Boolean rules among subsets of up to three input features selected from the available clinical features, e.g. {*A, B*, …, *L*}, see Fig. 1a.

**Fig. 1.**
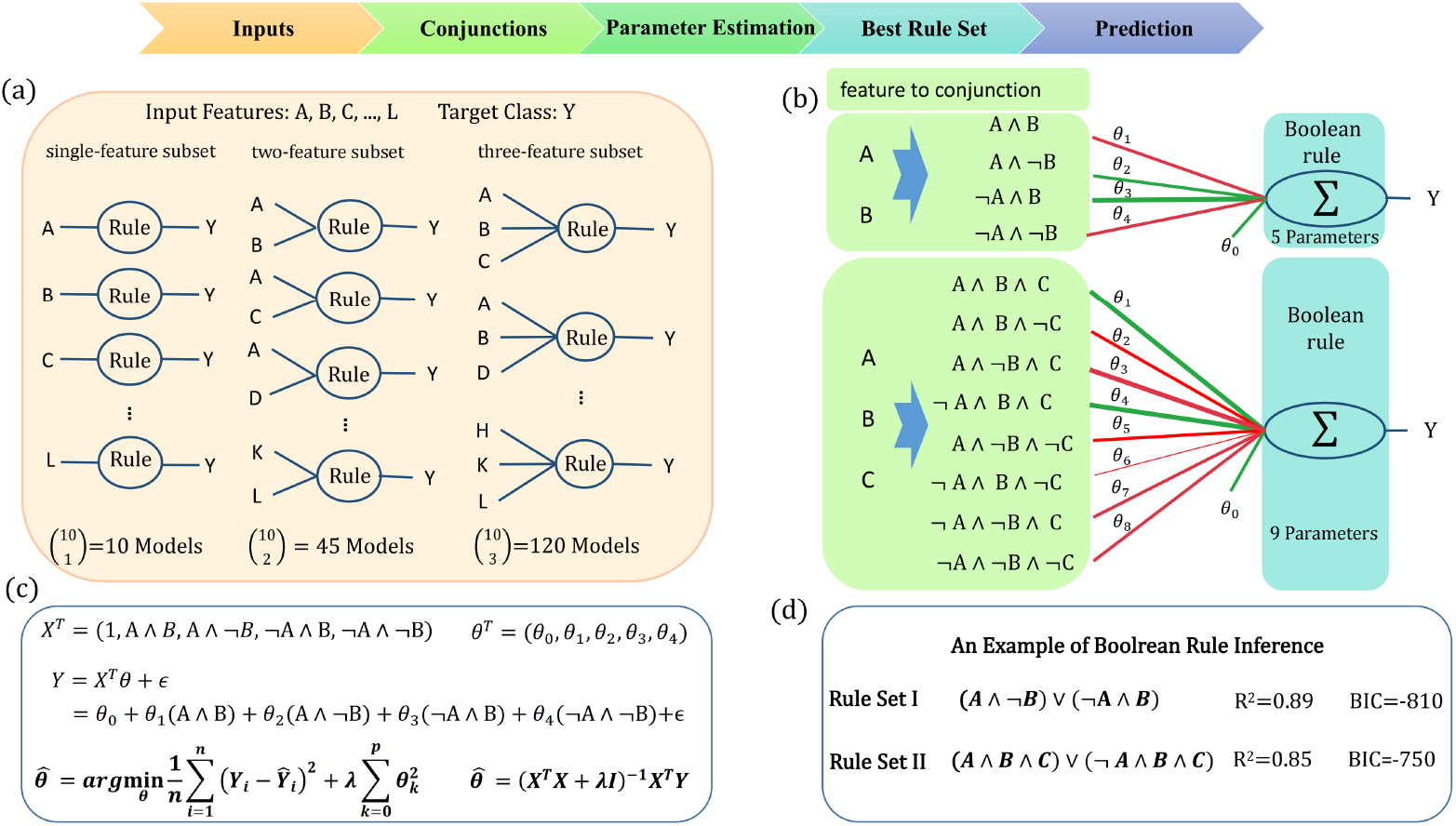
Illustration of Boolean rule inference in medical datasets with RBBR. (a) Ridge regression models are fitted to Boolean rule sets derived from single-feature, two-feature, and three-feature subsets of the input features *{A, B, C*, …, *H, K, L}* to predict patient outcomes. In this example with 10 input features, the number of single-feature, two-feature, and three-feature subsets are 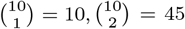, and 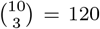, respectively. A distinct ridge regression model is fitted to the Boolean rules (conjunctions) constructed from each subset, with these rules serving as inputs and patient outcomes as the target. (b) Examples of Boolean rule inference from ridge regression models fitted to a two-feature subset including {*A, B*} and a three-feature subset including {*A, B, C*}. In each case, the input features are first transformed into Boolean rules or conjunctions. For example, the feature set {*A, B*} is transformed into the Boolean rules (*A* ∧ *B*), (*A* ∧ ¬*B*), (¬*A* ∧ *B*), and (¬*A* ∧ ¬*B*), which serve as inputs to predict the patient outcome in the ridge regression model. Boolean rules receiving positive coefficients are generally positively correlated with the patient outcome, e.g., they define conditions ensuring a value of 1 for the binary (0/1) target class. For a two-feature subset {*A, B*}, as in this example, the Boolean rules (*A*∧ ¬*B*) and (¬*A*∧ *B*) may receive positive coefficients (in green) in the fitted ridge regression, while other rules receive negative coefficients (in red). The outputs from all conjunctions are summed, considering their coefficients, to predict the target class. (c) Shows the ridge regression optimization framework, including the 𝓁_2_-regularization term, to estimate the rule coefficients. (d) Examples of Boolean rule sets (*A* ∧ ¬*B*) ∨ (¬*A* ∧ *B*) and (*A* ∧ *B* ∧ *C*) ∨ (¬*A* ∧ *B* ∧ *C*) inferred from a two-feature and a three-feature subset in panel (b), sorted based on the BIC of the associated regression model. ∧ denotes the Boolean AND operator, ∨ denotes the Boolean OR operator, and ¬ represents negation.

- **Single-feature subsets**: For a single-feature subset like {*A*}, the possible Boolean rules are {*A*, ¬*A*}.
- **Two-feature subsets**: For a two-feature subset like {*A, B*}, the possible Boolean rules are {(*A* ∧ *B*), (*A* ∧ ¬*B*), (¬*A* ∧ *B*), (¬*A* ∧ ¬*B*)}.
- **Three-feature subsets**: For a three-feature subset like {*A, B, C*}, the possible Boolean rules are {(*A* ∧ *B* ∧ *C*), (*A* ∧ *B* ∧ ¬*C*), …, (¬*A* ∧ ¬*B* ∧ ¬*C*)}.

Considering Boolean rules among up to three features ensures their interpretability for clinicians while maintaining reliable predictive power [13, 14]. Additionally, this constraint makes RBBR computationally efficient and applicable to high-dimensional datasets with many clinical features, as it focuses only on Boolean rules derived from small feature subsets that effectively capture variations in the target class. With this aim, in RBBR, a distinct ridge regression model is fitted to each Boolean rule set derived from a unique subset of input features. See Fig. 1b for an example of a regression model fitted to the rule set {(*A* ∧ *B*),(*A* ∧ ¬*B*),(¬*A* ∧ *B*),(¬*A* ∧ ¬*B*)} as input, with the target class as the response variable. Fig. 1b also illustrates a regression model fitted to the rule set {(*A*∧*B* ∧ *C*),(*A*∧*B* ∧ ¬*C*), …, (¬*A*∧ ¬*B* ∧ ¬*C*)} as input, again with the target class as the response variable. See Fig. 1c for the calculations used to estimate rule coefficients. To achieve an optimal balance between goodness of fit and model complexity (Boolean rule set size), optimal rules for explaining the target class are derived from the regression model that achieves the lowest Bayesian Information Criterion (BIC). For each ridge regression, the BIC is calculated as follows:

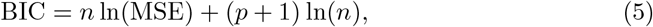

where 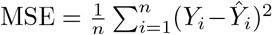 denotes the Mean Squared Error for a fitted regression model, *p* + 1 is the number of parameters (for *p* Boolean rules plus intercept), and *n* is the sample size. We utilize the BIC to rank distinct ridge regression models fitted to Boolean rule sets, each involving different subsets of input features. Among all fitted ridge regression models, the preferred model is the one with a lower BIC value [20], guiding the selection of Boolean rules with positive coefficients for predictive purposes and clinical diagnosis. We retain Boolean rules with positive coefficients, as they are likely positively correlated with the target class and define conditions ensuring a value of 1 for the binary (0/1) target class. Conversely, Boolean rules with negative coefficients in the fitted ridge regression model define conditions ensuring a value of regression model define conditions ensuring a value of 0. In Fig. 1b, Boolean rules with positive coefficients, are shown in green, while those with non-positive coefficients are shown in red. In Fig. 1b, for the Boolean rule set {(*A* ∧ *B*),(*A* ∧ ¬*B*),(¬*A* ∧ *B*),(¬*A* ∧ ¬*B*)} involving the two-feature subset {*A, B*}, rules like (*A* ∧ ¬*B*) or (¬*A* ∧ *B*) may receive positive coefficients, as in this example. This results in the combination (*A*∧¬*B*)∨(¬*A*∧*B*), which can be used to explain the target class *Y*. For a binary (0/1) target class, such rules would yield an outcome of 1, which could represent disease states or other clinical outcomes. Another example in Fig. 1b is a regression model fitted to the rule set {(*A*∧*B*∧*C*),(*A*∧*B*∧¬*C*), …, (¬*A*∧ ¬*B* ∧ ¬*C*)}, which involves the three-feature subset {A, B, C}. In this example, two rules, (*A* ∧ *B* ∧ *C*) ∨ (¬*A* ∧ *B* ∧ *C*), receive positive coefficients, indicating that these rules would yield an outcome of 1 for a binary (0/1) target class. Fig. 1d presents two predicted rule sets, (*A* ∧ ¬*B*) ∨ (¬*A* ∧ *B*) and (*A* ∧ *B* ∧ *C*) ∨ (¬*A* ∧ *B* ∧ *C*), inferred from the example ridge regression models in panel b. These rule sets are sorted based on the BIC values of the associated regression models.

### RBBR Evaluation and Complexity Metrics for Boolean Rules

The RBBR’s performance was assessed across six medical datasets, as presented in Table 1. The inferred Boolean rules with positive coefficients (*θ*_*j*_ *>* 0) were analyzed for clinical relevance, ensuring alignment with known disease pathways. Additionally, we plotted the Receiver Operating Characteristic (ROC) and Precision-Recall (PR) curves using 5-fold cross-validation across different datasets. The RBBR’s performance was benchmarked against other rule-based tools based on (i) accuracy metrics such as Accuracy (ACC) and Area Under the Receiver Operating Characteristic curve (AUROC) [21], and (ii) interpretability metrics, including Rule Number (RN), which represents the number of Boolean rules with positive coefficients (e.g., *RN* = 2 for (*A* ∧ ¬*B*) ∨ (¬*A* ∧ *B*)) in the fitted model, and Rule Length (RL), which indicates the number of features in a rule (e.g., *RL* = 1 for (*A*), *RL* = 3 for (*A* ∧ *B* ∧ *C*).

## Results and Applications

In this section, we apply RBBR to six medical datasets – *Lung Cancer Prediction, Three-level Lung Cancer, Wisconsin Breast Cancer, Diagnostic Breast Cancer, Heart Failure Prediction*, and *Early Stage Diabetes Risk* – demonstrating its ability to infer interpretable Boolean rules that elucidate relationships between clinical risk factors and patient outcomes. For each dataset, (i) we describe the key features and risk factors along with their clinical relevance, and (ii) we infer Boolean rules using RBBR to capture risk factor combinations that predict disease risk, with detailed analyses provided below for the lung cancer datasets.

### Boolean Rule Inference for Lung Cancer Datasets

RBBR is applied for Boolean rule inference to predict lung cancer outcomes across two distinct datasets: *Lung Cancer Prediction* and *Three-level Lung Cancer. Lung Cancer Prediction* dataset is available from Kaggle and includes 59 samples with 4 features: smoking status, age, AreaQ (interpreted as environmental exposure, likely air quality or pollution levels), and alcohol use. The target class is binary, distinguishing between the presence and absence of lung cancer, aligning with early detection efforts. The small sample size (59 samples) and limited features suggest a focus on core risk factors, offering simplicity for interpretable modeling. The *Three-Level Lung Cancer* dataset, also available from Kaggle, consists of 1,000 samples with 23 features. These features include demographic variables (e.g., age, gender), lifestyle factors (e.g., smoking, alcohol use), environmental exposures (e.g., air pollution, occupational hazards), and symptomatic indicators (e.g., fatigue, chest pain, swallowing difficulty). The target class represents a three-level lung cancer diagnosis – potentially mild, moderate, or severe – making it suitable for multi-class classification tasks. This dataset’s large sample size and diverse feature set provide a robust foundation for uncovering complex interactions among risk factors and symptoms, enabling detailed risk stratification for advanced lung cancer stages [22]. Lung cancer, a leading cause of mortality globally, often presents late, necessitating early detection tools [23]. Both datasets address this challenge, but their differences – in size, features, and targets – reflect distinct research goals: the first for screening, and the second for staging.

In the *Lung Cancer Prediction* dataset, Table 2 presents three Boolean rule sets inferred by RBBR from regression models with the optimal BIC values. Rule Set 1 involves the combination of risk factors Age, AreaQ, and Alcohol, unexpectedly omitting Smoking. The highest coefficients (0.94 and 1.09) are assigned to the Boolean rules (Age ∧ AreaQ ∧ Alcohol) and (Age ∧ ¬AreaQ ∧ ¬Alcohol), where the latter identifies a subpopulation of older patients who lack both AreaQ and Alcohol. See Table 2 for additional rules in Rule Set 1 that received lower coefficients. As described earlier, to infer such a rule set, the RBBR framework initially transformed Age, AreaQ, and Alcohol into eight Boolean rules or conjunctions, such as (Age ∧ AreaQ ∧ Alcohol), (Age ∧ AreaQ ∧ ¬ Alcohol), …, (¬Age ∧ AreaQ ∧ ¬ Alcohol). These rules serve as inputs into the ridge regression model to predict disease states, such as lung cancer in this example. For this three-feature subset, Fig. 2a shows the Mean Squared Error (MSE) versus the logarithm of *λ*, indicating that the MSE is minimized when log(*λ*) is approximately −2. In Fig. 2b, the rule coefficients estimated in the ridge regression model are plotted against the log(*λ*), with the coefficients corresponding to the optimal *λ* indicated by a vertical dashed line, in red. In Table 2, Rule Set 2 simplifies the risk factors to the Boolean rule (Age ∨ Alcohol), indicating that Alcohol use in combination with Age is a major risk factor for lung cancer prediction. Rule Set 3 incorporates Boolean rules such as (Age ∧ Smokes ∧ Alcohol) and (Age ∧ ¬ Smokes ∧ Alcohol), with coefficients of 1.22 and 0.75, respectively. This rule set also identifies other combinations of Age, Smokes, and Alcohol with lower coefficients in predicting cancer risk. For practical applications, clinicians should integrate these rule sets with imaging data and patient histories to achieve comprehensive diagnoses [24]. As presented in Table 2, Boolean Rule Set 1, with an *R*^2^ of 0.92 and a BIC of −207, offers the best predictive power and explainability for the target class compared to Rule Sets 2 and 3.

**Fig. 2.**
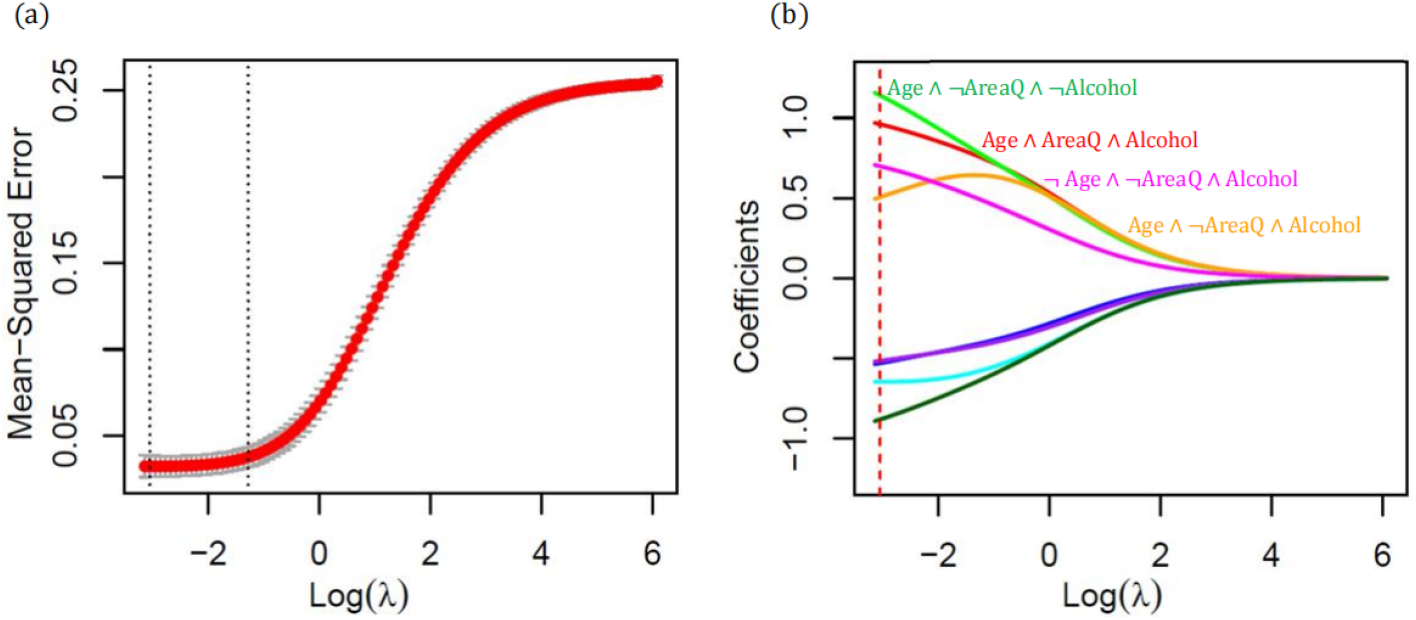
Estimation of Boolean rule coefficients in RBBR. For a three-feature subset involving Age, AreaQ, Alcohol in the Lung Cancer Prediction dataset, the Mean Squared Error (MSE) and estimated rule coefficients in the ridge regression model are plotted against log(*λ*). (a) The MSE is minimized for log(*λ*) *≈ −*2. (b) The rule coefficients for the optimal log(*λ*) are indicated with a vertical dashed line, in red.

**Table 2.**
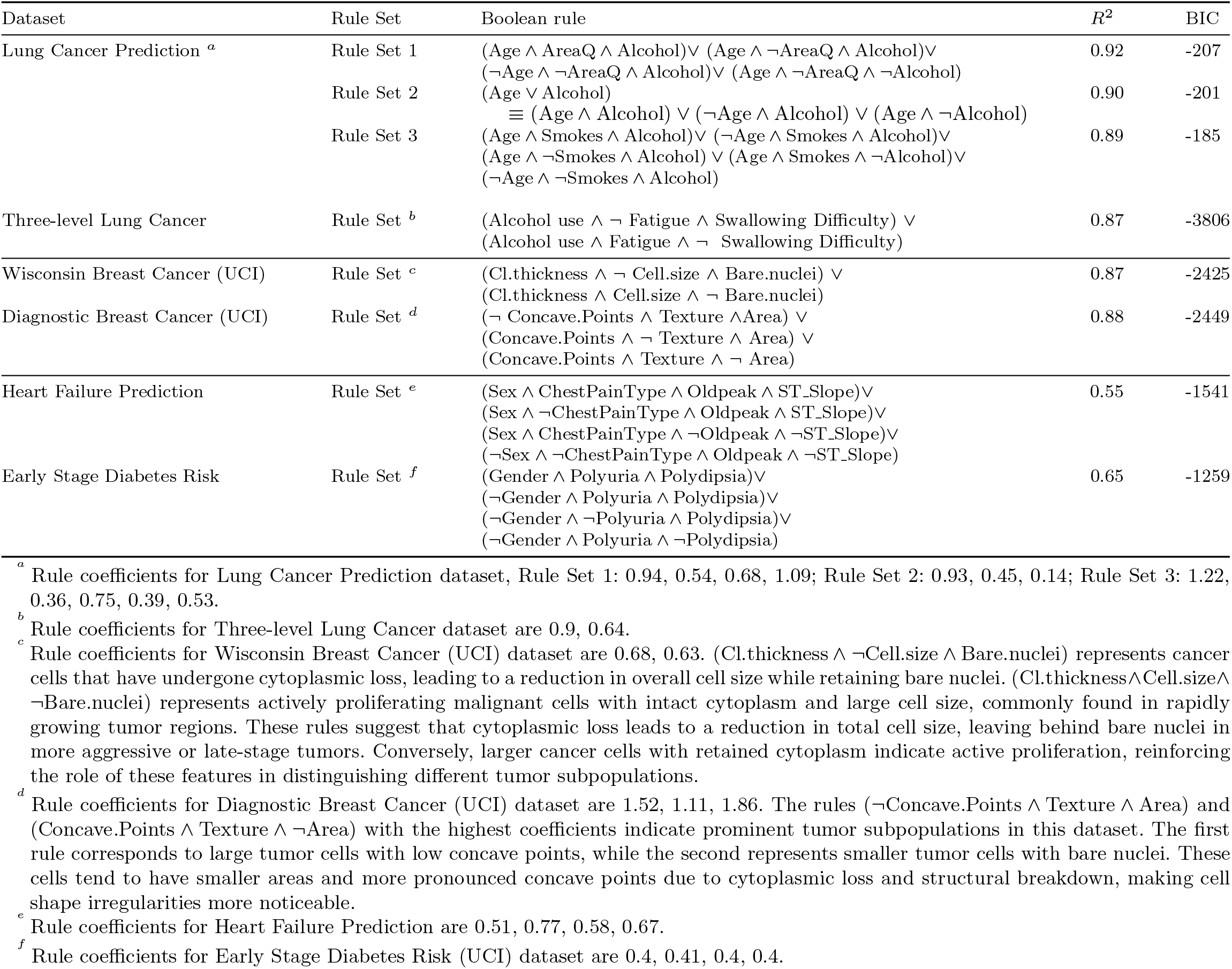
Boolean rules inferred by RBBR across the Lung Cancer, Breast Cancer, Heart Failure Prediction, and Early Stage Diabetes Risk datasets. *∧* denotes the Boolean AND operator, *∨* denotes the Boolean OR operator, and *¬* represents negation.

For the *Three-level Lung Cancer* dataset, the inferred rules highlight alcohol use, fatigue, and swallowing difficulty as major risk factors. The rule coefficients, presented in the footnote of Table 2, indicate the significance of these risk factor combinations in predicting lung cancer. Specifically, the rules (Alcohol use ∧ ¬ Fatigue ∧ Swallowing Difficulty) and (Alcohol use ∧ Fatigue ∧ ¬ Swallowing Difficulty) receive coefficients of 0.9 and 0.64, respectively, suggesting that alcohol use, in combination with either swallowing difficulty or fatigue, is a key risk factor for lung cancer diagnosis. Additionally, the rule (Alcohol use ∧ ¬ Fatigue ∧ Swallowing Difficulty) receives a higher coefficient than (Alcohol use ∧ Fatigue ∧ ¬ Swallowing Difficulty), indicating that patients with alcohol use, swallowing difficulty, and no fatigue may face a higher risk. This suggests that such a combination of risk factors is more prevalent among lung cancer patients, whereas patients with alcohol use, fatigue, and no swallowing difficulty may be less common in the dataset. In both lung cancer datasets, the Boolean rule sets inferred by RBBR distill complex feature interactions into interpretable rules that emphasize key symptoms and risk factors, aligning with clinical needs such as identifying high-risk patients for low-dose CT screening or symptom-based triage [24].

### Boolean Rule Inference for Breast Cancer Datasets

RBBR is applied for Boolean rule inference to predict breast cancer across two distinct datasets: *Wisconsin Breast Cancer* and *Diagnostic Breast Cancer*. These datasets are widely used in machine learning for medical diagnostics and provide rich feature sets, offering an opportunity to explore RBBR’s capabilities in inferring Boolean rules. The *Wisconsin Breast Cancer* (UCI) dataset comprises 699 samples with 9 features extracted from fine needle aspiration (FNA) cytology, including clump thickness, cell size, and bare nuclei. It is designed for binary classification to differentiate between benign and malignant tumors and serves as a benchmark in breast cancer research [25]. The *Diagnostic Breast Cancer* (UCI) dataset consists of 569 samples and 30 features, including measurements such as texture, area, and concave points derived from digitized FNA images. Like the Wisconsin dataset, it supports binary classification and is valued for its detailed imaging-derived features [26]. Table 2 presents the Boolean rules inferred by RBBR for these datasets, along with their corresponding *R*^2^ and BIC values. For the *Wisconsin Breast Cancer* (UCI) dataset, the inferred Boolean rules (Cl.thickness ∧ ¬Cell.size ∧ Bare.nuclei) and (Cl.thickness ∧ Cell.size ∧ ¬Bare.nuclei) receive the highest coefficients of 0.68 and 0.63, respectively. A few other rules with lower coefficients are also identified. These results suggest that Clump.thickness, in combination with either Cell.size or Bare.nuclei, plays a key role in distinguishing between benign and malignant breast cancers in this classification task. Clinically, such risk factors are justifiable, as malignant tumors tend to have higher clump thickness due to uncontrolled cell growth and the loss of normal cell organization. Additionally, larger cells suggest rapid, uncontrolled growth, which is a hallmark of cancer [27]. Furthermore, cells with bare nuclei (nuclei lacking surrounding cytoplasm) are prevalent in aggressive tumors due to cellular breakdown or death. Such cells are often observed in apoptotic or necrotic tumor regions, where rapid proliferation and insufficient oxygenation contribute to cellular breakdown. Intriguingly, the rule set (Cl.thickness∧ ¬Cell.size∧ Bare.nucle) ∨ (Cl.thickness∧ Cell.size∧ ¬Bare.nucle) suggests that malignant tumor cells are characterized either by bare nuclei or a large cell size. This implies that as malignant cells undergo surface membrane breakdown and cytoplasmic loss, initially large cancerous cells may be reduced to bare nuclei, a feature often observed in aggressive tumors. With an *R*^2^ of 0.87 and a BIC of −2425, the inferred rule set closely mirrors pathologist assessments, making it a reliable diagnostic aid. For instance, such rules could serve as decision-support tools in pathology labs, complementing expert review by flagging cases with high clump thickness, large cell sizes, and bare nuclei for urgent attention, as demonstrated by AI-driven diagnostic tools in dermatology [28].

For the *Diagnostic Breast Cancer* (UCI) dataset, the inferred Boolean rules (¬Concave.Points ∧ Texture ∧ Area), (Concave.Points ∧ ¬Texture ∧ Area), and (Concave.Points ∧ Texture ∧ ¬ Area) receive the highest coefficients of 1.52, 1.11, and 1.86, respectively. These rules suggest that a combinatorial assessment of Concave Points, Texture, and Area is effective for breast cancer diagnosis in this dataset. This aligns with clinical practice, as a large cell area in FNA images indicates a significant tumor mass, often associated with advanced disease [29]. Additionally, texture – a measure of local intensity variation – reflects heterogeneity within the tumor, with increased texture irregularity often linked to malignancy [30]. Numerous concave points – irregular, jagged cell edges – are characteristic of malignant cells with spiculated shapes [2]. The inferred Boolean rule sets in the Wisconsin and Diagnostic Breast Cancer datasets align closely with clinical expectations. These rules leverage combinatorial interactions among features – such as clump thickness, texture, cell size, area, and shape characteristics (e.g., concave points) – which reflect the microscopic and imaging signatures of breast cancer. Their interpretability and clinical relevance make them valuable tools for guiding diagnosis in real-world settings.

### Boolean Rule Inference for Heart and Diabetes Datasets

RBBR is additionally applied for Boolean rule inference in two medical datasets: the *Heart Failure Prediction* and *Early Stage Diabetes Risk* datasets. The *Heart Failure Prediction* dataset is sourced from the UCI Cleveland Heart Disease dataset and consists of 746 patient records with 11 clinical and demographic features, including age, sex, chest pain type, resting blood pressure, serum cholesterol, and exercise test outcomes (e.g., Oldpeak, ST Slope). Designed for binary classification, this dataset predicts the presence or absence of heart disease using clinical and stress testing features, with the goal of developing predictive models for early cardiovascular disease detection and improving patient outcomes through timely intervention [17]. The *Early Stage Diabetes Risk* dataset is also sourced from the UCI Machine Learning Repository. It includes 520 patient records with 16 features, such as age, gender, and symptoms like polyuria (excessive urination) and polydipsia (excessive thirst). This binary classification task aims to identify early-stage diabetes risk factors [17] based on demographic and clinical features, supporting preventive healthcare strategies.

For the *Heart Failure Prediction* dataset, the Boolean rule set in Table 2 suggests that, in males, the presence of either a specific chest pain type or significant ST depression (Oldpeak), along with a flat or downsloping ST segment (ST Slope), indicates heart disease. In contrast, for females, risk is flagged when Oldpeak is present while other clinical indicators are absent or reversed (e.g., an upsloping ST segment), reflecting known gender differences in cardiovascular failure. With an *R*^2^ of 0.55, this rule set captures critical ischemic indicators that can guide further diagnostic evaluation, such as coronary angiography [31].

For the *Early Stage Diabetes Risk* dataset, the Boolean rules inferred by RBBR indicate that for males, risk is predicted only when both polyuria and polydipsia are present (Gender ∧ Polyuria ∧ Polydipsia), reflecting the classical diabetic state driven by hyperglycemia-induced osmotic diuresis. In contrast, for females, the threshold is broader, with risk indicated even when only one of these symptoms is present (¬Gender ∧ Polyuria ∧ Polydipsia) ∨ (¬Gender ∧ ¬Polyuria ∧ Polydipsia) ∨ (¬Gender ∧ Polyuria ∧ ¬ Polydipsia). This difference may reflect physiological variations in symptom manifestation but also raises concerns about potential over-diagnosis in females [32]. With an *R*^2^ of 0.65, this Boolean rule set provides a rational basis for early risk screening, guiding the need for confirmatory tests such as fasting glucose or HbA1c. In both the heart failure and diabetes datasets, Boolean rules capture gender-specific risk factors and subtle physiological differences between genders that align with established clinical knowledge. These findings demonstrate that RBBR infers clinically meaningful rules that can serve as preliminary risk indicators, guiding further diagnostic workup.

### ROC and PR Curve Analysis for RBBR Performance

In Fig. 3, RBBR is further evaluated by plotting ROC and PR curves across five medical datasets with a binary target class. The ROC curves, which plot sensitivity against the false positive rate (FPR), show that RBBR achieves high area under the curve values – close to or above 0.9 for most datasets, such as *Lung Cancer Prediction* (red), *Wisconsin Breast Cancer* (green), and *Diagnostic Breast Cancer* (blue) – indicating excellent discriminative ability in distinguishing diseased from non-diseased cases [33]. Similarly, the PR curves, which plot precision against recall, highlight RBBR’s effectiveness in handling imbalanced medical datasets, with curves maintaining high precision even at high recall levels. These metrics highlight RBBR’s reliability in delivering accurate predictions, complementing its previously reported *R*^2^ and BIC values, and reinforcing its value for clinical decision-making across diverse medical contexts [22].

**Fig. 3.**
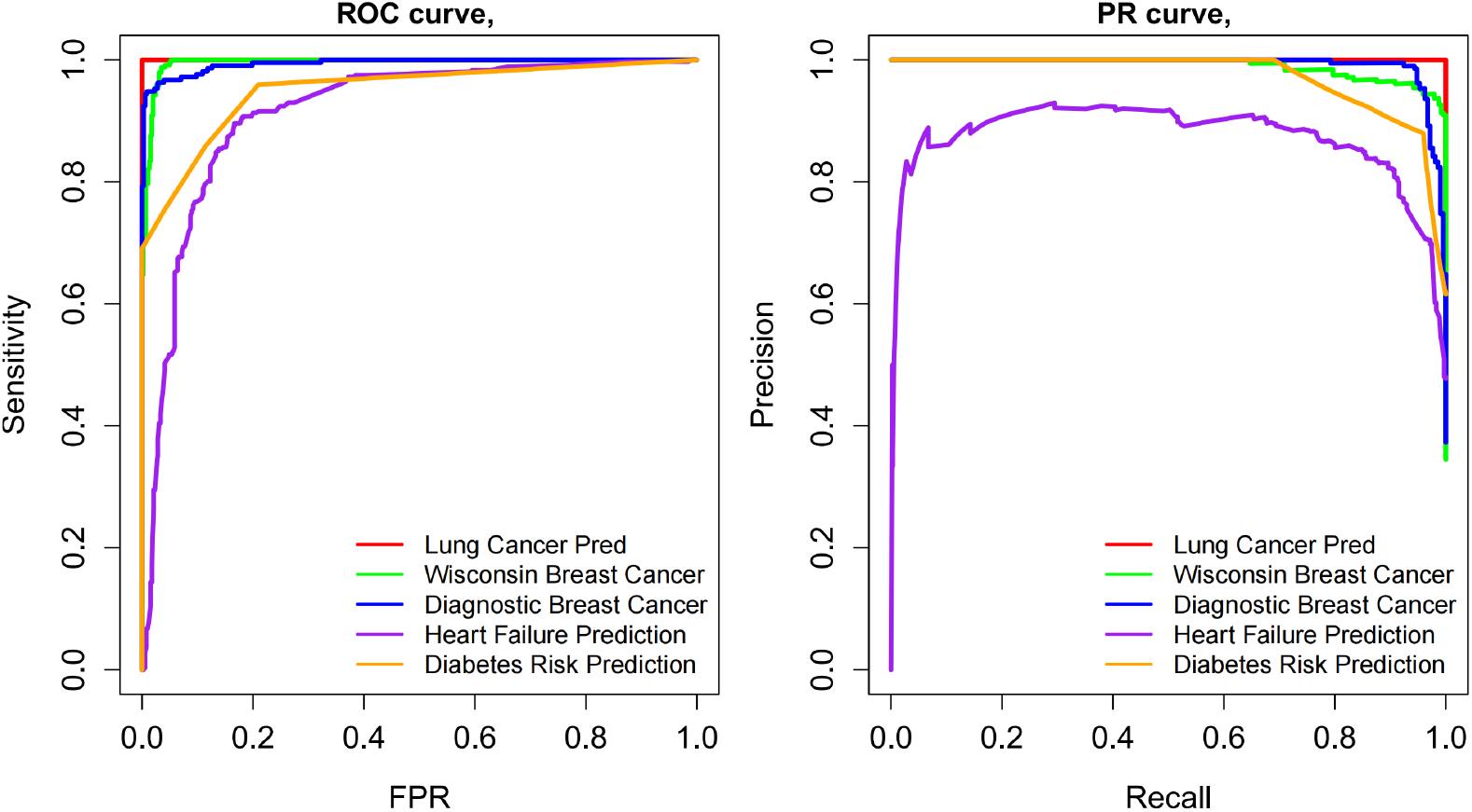
ROC and PR curves are plotted to evaluate RBBR performance across five medical datasets with a binary target class.

### Benchmarking RBBR Performance Against Other Rule-Based Methods

The performance of RBBR is further benchmarked against three prominent rule-based tools – DeepRED, REM-D, and ECLAIRE [34]. These tools were developed to enhance interpretability in DNNs by inferring rule sets from trained weights in DNNs. To achieve this, they rely on decision tree algorithms, such as C5.0, to map the hidden layers of DNNs to the target class. In Table 3, the performance of RBBR is compared to DeepRED, REM-D, and ECLAIRE across two medical datasets: *Lung Cancer Prediction* and *Diagnostic Breast Cancer*. Each tool is evaluated using metrics such as Accuracy (ACC) and AUROC for predictive performance, along with Rule Number (RN) and Rule Length (RL) to assess interpretability and rule complexity. The mean and standard deviation of each metric are reported across 5-fold cross-validation for each dataset. Bold values indicate the best performance within each dataset.

**Table 3.** Benchmarking RBBR against rule-based tools (DeepRED, REM-D, and ECLAIRE) on two medical datasets. The average and standard deviation of accuracy metrics (ACC, AUROC) and rule complexity metrics (RN, RL) are reported over 5-fold cross-validation.

| Dataset | Method | ACC (%) | AUROC (%) | RN | RL |
| --- | --- | --- | --- | --- | --- |
| Lung Cancer Prediction | DeepRED | 59.5 ± 17.4 | 59.6 ± 17.1 | **2.0** ± 0.0 | **0.0** ± 0.0 |
| | REM-D | 59.5 ± 17.4 | 59.6 ± 17.1 | **2.0** ± 0.0 | **0.0** ± 0.0 |
| | ECLAIRE | 93.2 ± 3.4 | 93.6 ± 3.2 | 3.6 ± 1.86 | 1.18 ± 0.23 |
| | RBBR | **98.3** ± 3.7 | **98.0** ± 4.5 | 3.2 ± 0.45 | 2.2 ± 0.447 |
| Diagnostic Breast Cancer | DeepRED | 91.4 ± 1.5 | 89.9 ± 2.2 | **2.0** ± 0.0 | **1.0** ± 0.0 |
| | REM-D | 91.4 ± 1.5 | 90.0 ± 2.0 | **2.0** ± 0.0 | **1.0** ± 0.0 |
| | ECLAIRE | 94.4 ± 2.5 | 93.9 ± 2.7 | 32.8 ± 12.73 | 2.13 ± 0.27 |
| | RBBR | **96.5** ± 2.1 | **95.9** ± 2.3 | 4.6 ± 0.55 | 3 ± 0 |

RBBR consistently outperforms or closely matches the best-performing methods in ACC and AUROC. In the *Lung Cancer Prediction*, over 5-fold cross-validation, RBBR achieves an average ACC of 98.3% and AUROC of 98.0%, significantly surpassing DeepRED and REM-D (both with ACC = 59.5% and AUROC = 59.6%) and outperforming ECLAIRE (ACC = 93.2%, AUROC = 93.6%). In terms of interpretability metrics, the average RN and RL over 5-fold cross-validation for RBBR are 3.2 and 2.2, respectively, compared to 3.6 and 1.18 for ECLAIRE, the second-best performing tool in terms of ACC. DeepRED and REM-D produce the simplest rule sets (e.g., RN = 2 and RL = 0 for *Lung Cancer Prediction*), but their low accuracy limits clinical utility. In the *Diagnostic Breast Cancer*, RBBR leads with an average ACC of 96.5% and AUROC of 95.9%, outperforming ECLAIRE and the other methods. In terms of interpretability metrics, the average RN and RL over 5-fold cross-validation for RBBR are 4.6 and 3.0, respectively, compared to 32.8 and 2.13 for ECLAIRE, the second-best performing tool. Compared to ECLAIRE, RBBR relies on fewer rules for prediction, making it more practical for clinical applications.

## Conclusion

The Boolean rule sets inferred by RBBR serve as a checklist for clinicians, prioritizing key diagnostic features and enhancing confidence in verifying diagnoses. In breast cancer, the inferred rules – incorporating features such as cell area, size, irregular edges, clump thickness, texture, or bare nuclei – closely align with histopathological observations and imaging findings [27]. For *Heart Failure Prediction*, RBBR uncovered gender-specific differences in stress test results (e.g., variations in ST Slope and Oldpeak). In *Early Stage Diabetes Risk*, women appeared to signal risk with fewer symptoms (e.g., excessive thirst alone) compared to men, a finding consistent with established clinical disparities [35, 36].

A notable strength of RBBR is its interpretability, achieved through Boolean logic (e.g., if A ∧ B, then risk) combined with ridge regression for efficiently estimating rule coefficients. RBBR demonstrates strong predictive performance compared to rule-based tools such as DeepRED, REM-D, and ECLAIRE, while maintaining interpretability for clinical applications that require trust, transparency, and actionable decision-making. Each rule in a Boolean rule set inferred by RBBR could correspond to a specific class of tumor cells or disease subpopulations. For instance, the rule set (Cl.thickness∧¬Cell.size∧Bare.nuclei) ∨ (Cl.thickness∧Cell.size∧¬Bare.nuclei) derived from the *Wisconsin Breast Cancer* (UCI) dataset suggests that malignant cells are characterized either by the presence of bare nuclei or by a large cell size. This implies that initially large cancerous cells may shrink to bare nuclei as they undergo surface membrane breakdown and cytoplasmic loss, a phenomenon commonly observed in aggressive tumors. By inferring clinically relevant Boolean rules to reliably predict patient outcomes or disease status, RBBR provides actionable insights that support, rather than replace, clinical judgment – a pivotal step toward making medical AI more transparent and trustworthy. All datasets used in this study are publicly available from Kaggle (https://www.kaggle.com/) and the UCI Machine Learning Repository (https://archive.ics.uci.edu/)[17].

## Supplementary Information

None.

## Acknowledgements

None.

## Author contributions

M.E. performed the data curation, formal analysis, and validation. M.E. also contributed to the visualization, writing of the original draft, and review and editing of the manuscript. S.A.M. contributed to the conceptualization and methodology, and participated in the review and editing of the manuscript. All authors reviewed the manuscript.

## Data availability

The datasets used in this study are publicly available from Kaggle and the UCI Machine Learning Repository.

## Declarations

### Ethical approval

The study adheres to the Helsinki declaration.

### Consent for publication

There was no patient or public involvement in the design, conduct, reporting, interpretation, or dissemination of this study.

